# Derivation of Isogenic Mesodermal and Ectomesodermal Chondrocytes from Human Induced Pluripotent Stem Cells for Articular Cartilage Regeneration

**DOI:** 10.1101/2020.10.02.323857

**Authors:** Ming-Song Lee, Matthew J. Stebbins, Hongli Jiao, Hui-Ching Huang, Brian E. Walzack, Sean P. Palecek, Eric V. Shusta, Wan-Ju Li

## Abstract

Generating phenotypic chondrocytes from human pluripotent stem cells through driving developmental lineage-specific differentiation remains to be of great interest in the field of cartilage regeneration. In this study, we derived chondrocytes from human induced pluripotent stem cells (hiPSCs) along the mesodermal or ectomesodermal lineages to prepare isogenic mesodermal cell-derived chondrocytes (MC-Chs) or neural crest cell-derived chondrocytes (NCC-Chs), respectively, and further evaluated differences in their cellular and molecular characteristics and cartilage repair capabilities. Our results showed that both lineage-derived chondrocytes expressed hyaline cartilage-associated markers and were capable of forming hyaline cartilage-like tissue ectopically and at joint defects. Moreover, NCC-Chs showed the absence of markers of hypertrophic chondrocytes and revealed a closer morphological resemblance to articular chondrocytes and a greater capability of producing glycosaminoglycans and collagen type 2 at cartilage defects compared to MC-Chs. It was found that the profile of global transcript expression of NCC-Chs more closely resembled that of native chondrocytes (NCs) than that of MC-Chs. Induced by additional growth factors identified through the analysis of transcriptome comparison to NCs, both MC-Chs and NCC-Chs showed a further increase in the phenotype of hyaline cartilage chondrocytes. Results of this study reveal differences in cellular and molecular characteristics and cartilage repair capabilities between isogenic hiPSC-derived MC-Chs and NCC-Chs and demonstrate that chondrocytes derived from hiPSCs along the ectomesodermal lineage are a potential cell source for articular cartilage regeneration.

## Introduction

Hyaline cartilage, originating from either mesoderm or ectoderm, is a connective tissue on an articular surface to transmit loads or in the nasal septum to provide structural support (*1*). During development, the mesoderm layer develops into somite and lateral plates, and cells of these mesodermal structures further generate hyaline cartilage in the spine and limbs, while the ectoderm layer gives rise to external ectoderm, neural tube, and neural crest (*2*). Specifically, neural crest, occasionally called “the fourth germ layer” due to its astonishing multipotency, is able to generate a variety of ectodermal and mesodermal cell types. During craniofacial development, neural crest-derived cells (NCCs), as an outgrowth of epithelium cells near the neural tube (*3*), convert into ectomesodermal cells and then become cranial NCCs. This allows the formation of head mesenchyme for further generation of craniofacial hyaline cartilage (*4*). Since mesoderm-derived cells (MCs) and NCCs share the capacity to generate hyaline cartilage, their progeny is considered appropriate cell sources for hyaline cartilage regeneration.

Mesenchymal stem/stromal cells (MSCs) derived from adult tissues, such as bone marrow and fat, possess the multilineage differentiation ability and have been extensively studied as a promising cell source for tissue regeneration such as cartilage reconstruction. To date, there are more than a thousand ongoing clinical trials using MSCs for therapies (*5*). While many of the trials focus on cartilage repair, increasing evidence has suggested that MSCs might not be a suitable cell type as a therapeutic agent. In particular, results of a clinical study have shown fibrous cartilage formation in repaired joint defects implanted with MSCs (*6*). Another study has revealed that chondrogenesis of MSCs leads to the production of network-forming collagen type 10 alpha 1 chain (COL10A1), similar to that generated by hypertrophic chondrocytes in the growth plate (*7*). Considering that the mechanical property of both fibrous and hypertrophic cartilage is inferior to that of hyaline cartilage, it becomes less attractive to use MSCs for cartilage regeneration.

Human embryonic stem cells (hESCs) and human induced pluripotent stem cells (hiPSCs) share similar characteristics and differentiation potential (*8*), and are considered alternative stem cell sources for cartilage regeneration. Studies have demonstrated induction of pluripotent stem cells (PSCs) into chondrocytes along the mesodermal or ectomesodermal lineage. For example, several groups have shown that through stepwise induction by growth factors, mesoderm-derived chondrocytes can be generated from hESCs (*9, 10*) or hiPSCs (*11–14*), and the mesoderm-derived cell is capable of repairing cartilage lesions in rodent joints (*12, 13, 15*). Besides mesodermal derivation, other studies have shown that chondrocytes can be obtained through differentiation of NCCs derived from hESCs (*16–18*) or hiPSCs (*19*) along the ectomesodermal lineage. Recently, our group has also reported that chondrocytes differentiated from hiPSC-derived NCCs express cartilage-associated markers and matrix (*20*). These findings suggest that cartilage formation during development can be emulated, to some degree, by induction of *in vitro* chondrogenic differentiation of PSCs along distinct developmental lineages.

While previous studies indicate the feasibility of chondrogenic differentiation of PSCs through stepwise induction by chemically defined medium, it remains unclear whether isogenic mesodermal and ectomesodermal lineage-derived chondrocytes from hiPSCs are different in phenotypes and properties and how similar the derived cells are to native chondrocytes (NCs). In this study, blood-derived hiPSCs were induced for mesodermal and ectomesodermal differentiation to prepare isogenic MCs and NCCs, respectively, following modified published protocols (*9, 17*), and then induced to generate chondrocytes. Differences in phenotypes and capability of cartilage regeneration between the distinct lineage-derived chondrocytes were characterized by both *in vitro* and *in vivo* assays. Genome-wide transcriptome analysis was also performed to compare hiPSC-derived chondrocytes to NCs to detect critical differences and identify potential molecular candidates for priming chondrogenic induction of MCs and NCCs and enhancing hyaline cartilage formation.

## Results

### Blood-derived hiPSCs differentiate into chondrocyte-like cells along mesodermal or ectomesodermal lineages

The pluripotency of hiPSCs was examined by the analysis of cell morphology, characterization of pluripotency markers, and teratoma formation. The results showed that compact colonies similar to those of hESCs were formed in culture (fig. S1A). Immunofluorescence and flow cytometry results demonstrated that the colonies exhibited pluripotency-related markers, including alkaline phosphatase (ALP), OCT4, NANOG, SOX2, PODPCALYXIN, SSEA4, and CD9 (fig. S1B-D). Results of teratoma formation in severe combined immunodeficient (SCID) mice showed that the hiPSC was capable of giving rise to ectodermal, endodermal, and mesodermal lineage-derived cells (fig. S2). All these together confirm the pluripotency of blood-derived hiPSCs.

To determine phenotypic differences between chondrocytes derived from the two different developmental lineages, blood-derived hiPSCs were induced into MCs and NCCs and further into isogenic MC-derived chondrocytes (MC-Chs) and NCC-derived chondrocytes (NCC-Chs), respectively (Fig. 1A), following previously published protocols with modifications (*9, 20*). The morphology of mesendodermal cells showed similar compactness to that of hiPSCs at day 4, while the morphology of differentiated mesoderm revealed cell cluster formations at day 8 (Fig. 1B). On the other hand, during induction, NCCs exhibited compact morphology at day 5 and later became stellate-shaped at day 15 (Fig. 1B). Results of immunofluorescence assessments indicated that MCs expressed mesoderm markers, CD34 and alpha-smooth muscle actin (a-SMA), after 8 days of MC induction (Fig. 1C), and NCCs were stained positive for neural crest markers, P75 and HNK1, at day 15 of induction (Fig. 1C). These results demonstrate that the derivation of MCs and NCCs from hiPSCs was achieved with the use of the induction protocols.

**Fig. 1.**
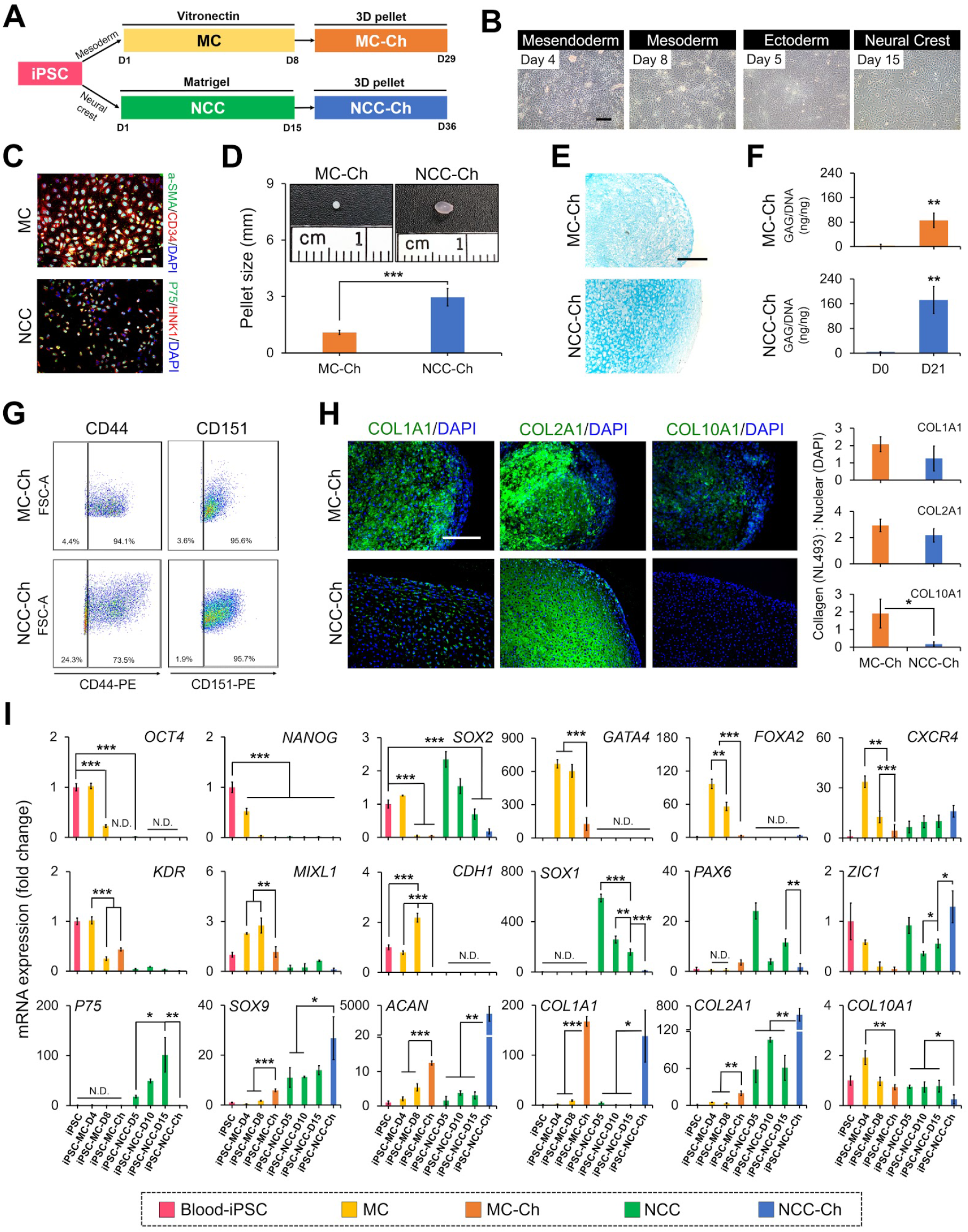
Differentiation of blood-derived hiPSCs toward isogenic MC-Chs and NCC-Chs. **(A)** Schematic of procedures inducing differentiation from hiPSCs to chondrocytes and corresponding timelines. hiPSCs were induced to differentiate into MCs and NCCs and then into MC-Chs and NCC-Chs, respectively. **(B)** Morphology of mesodermal and ectomesodermal lineage cells determined by microscopic imaging. **(C)** Immunofluorescence staining of hiPSC-derived MCs for detection of a-SMA and CD34 at day 8 and that of hiPSC-derived NCCs for detection of P75 and HNK1 at day 15. Nuclear DNA was labeled by DAPI. **(D)** Representative macrographs and size quantification of MC-Ch and NCC-Ch pellets. **(E)** MC-Ch and NCC-Ch pellets stained by Alcian blue. **(F)** Quantification of GAG in MC-Ch and NCC-Ch pellets analyzed by dimethylmethylene blue. **(G)** Flow cytometry analysis of cells for detection of chondrocyte-related markers, CD44 and CD151. **(H)** Immunofluorescence staining of MC-Ch and NCC-Ch pellets for detection of COL1A1, COL2A2, and COL10A1. Nuclear DNA was labeled with DAPI. **(I)** Dynamics of the mRNA expression of pluripotency markers (*OCT4, NANOG*, and *SOX2*), mesoderm-associated markers (*GATA4, FOXA2, CXCR4, KDR, MIXL1*, and *CDH1*), neural crest-associated markers (*SOX1, PAX6, ZIC1*, and *P75*), and cartilage-associated markers (*SOX9, ACAN, COL1A1, COL2A1*, and *COL10A1*) during differentiation of hiPSCs into chondrocytes. Bars are color-coded to represent different cell types. *n* = 3. \**p* < 0.05, \*\**p* < 0.01, \*\*\**p* < 0.001. Scale bar, 200 μm.

Specific lineage-derived chondrocytes were generated through chondrogenic induction of cell pellets, and the results showed that pellets of NCC-Chs were significantly larger than those of MC-Chs (Fig. 1D). NCC-Ch pellets demonstrated stronger staining of Alcian blue (Fig. 1E) and higher glycosaminoglycan (GAG) content (Fig. 1F) than MC-Ch ones. The flow cytometry result showed that chondrocyte-related surface markers, CD44 and CD151, were expressed in 94.1% and 95.6% of MC-Chs and in 73.5% and 95.7% of NCC-Chs, respectively (Fig. 1G). Immunofluorescence staining indicated that the expression of collagen type 1 alpha 1 chain (COL1A1) and collagen type 2 alpha 1 chain (COL2A1) in MC-Ch pellets was comparable to that in NCC-Ch pellets while the COL10A1 content of MC-Ch pellets was greater than that of NCC-Ch pellets (Fig. 1H). Since COL10A1 is a marker of hypertrophic chondrocytes (*21*), the results suggest that the cells of MC-Ch pellets undergo hypertrophy.

To identify cell derivatives at different stages of differentiation from hiPSCs to NCC-Chs or MC-Chs, qPCR was performed to detect the mRNA expression of cell specific markers (Fig. 1I). The expression of pluripotency markers, *OCT4, NANOG,* and *SOX2,* highly present in hiPSCs gradually decreased during differentiation and was extremely low in MC-Chs and NCC-Chs. The mesoderm-related markers, *GATA4, FOXA2,* CXCR4, *KDR, MIXL1,* and *CDH1*, and the neural crest-related markers, *SOX1, PAX6, ZIC1*, and *P75*, were highly expressed in MCs and NCCs, respectively. The progressive increase in the expression of cartilage-associated markers, *SOX9* and *ACAN,* during chondrogenesis of both lineage cells indicated successful generation of MC-Chs and NCC-Chs from hiPSCs. Other cartilage associated markers, *COL1A1* and *COL2A1,* significantly increased, whereas *COL10A1* markedly decreased, in both lineage-derived chondrocytes compared to those in MCs and NCCs. Altogether, these results indicate that MC-Chs and NCC-Chs derived from blood-derived hiPSCs are chondrocyte-like cells expressing hyaline cartilage-associated extracellular matrix (ECM) and markers.

### MC-Ch and NCC-Ch implants form hyaline cartilage-like tissue ectopically in mice

Pellets of MC-Chs and NCC-Chs after 21 days of chondrogenic induction were subcutaneously implanted in mice for 30 days to ectopically generate hyaline cartilage. Semi-transparent and smooth MC-Ch and NCC-Ch pellets significantly increased from around 1 to 4 mm and 4 to 7 mm in size, respectively (Fig. 2A). The increase in pellet size is likely resulted from the accumulation of cartilage-associated ECM. The morphology of MC-Chs stained by haematoxylin and eosin (H&E) appeared to be spindle-shaped while that of NCC-Chs was round-shaped (Fig. 2B), suggesting that, compared to MC-Chs, NCC-Chs in lacunae more closely resemble spherical chondrocytes in native hyaline cartilage. Safranin O staining revealed that GAG-rich matrix was accumulated in MC-Ch and NCC-Ch pellets (Fig. 2B). The immunofluorescence analysis detecting human vimentin showed that the harvested pellets comprised mainly of human cells (Fig. 2C). Moreover, both MC-Ch and NCC-Ch pellets contained abundant COL2A1 and a small amount of COL1A1 while there was significantly more COL2A1 and less COL1A1 produced by NCC-Chs than that by MC-Chs (Fig. 2D). COL10A1 was sparsely present in MC-Ch pellets, whereas the molecule was absent in NCC-Ch pellets. Taken together, these results suggest that both chondrocyte lines form cartilage-like tissue subcutaneously in mice, but the tissue derived from a NCC-Ch pellet is more similar to hyaline cartilage than that from a MC-Ch pellet.

**Fig. 2.**
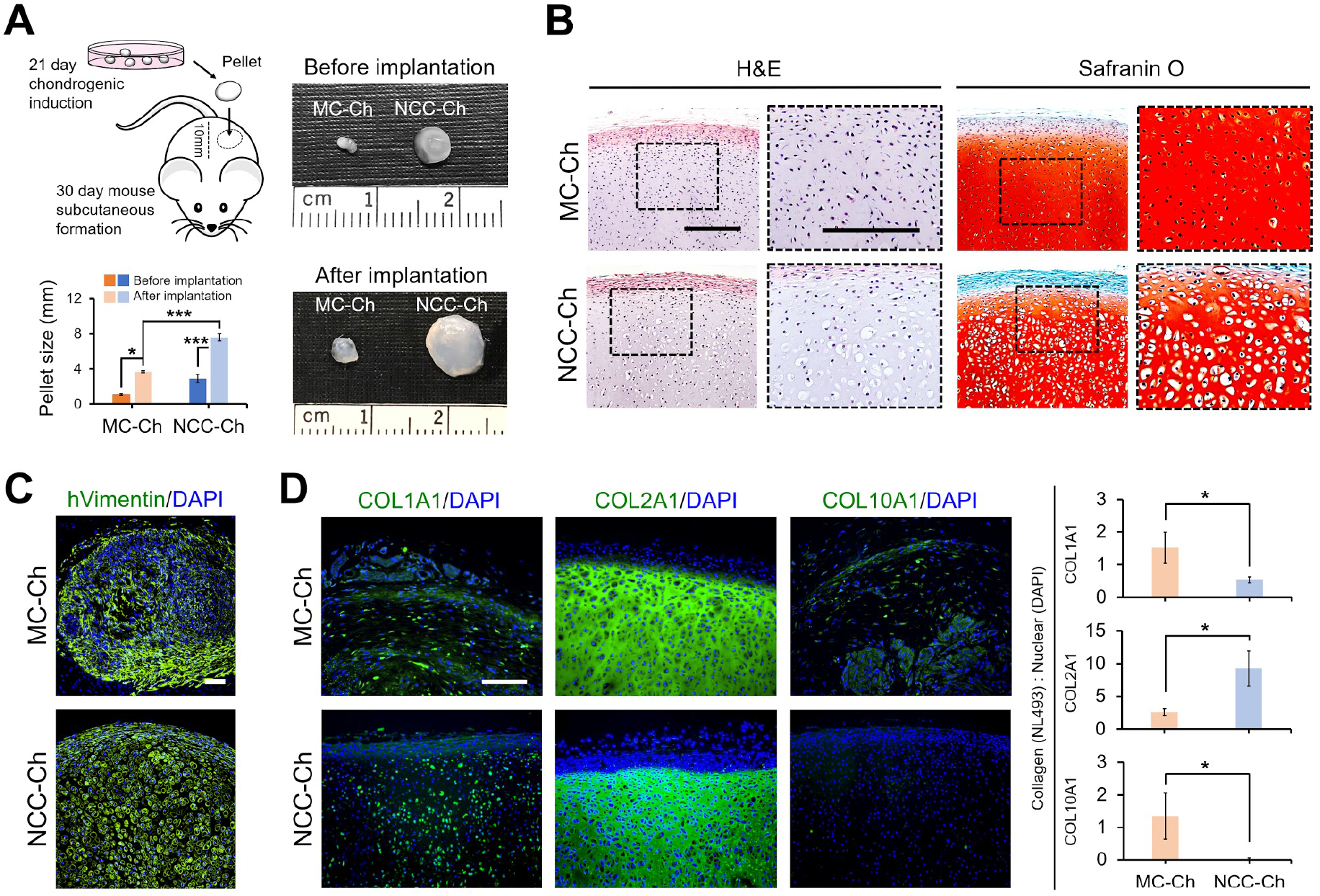
Subcutaneous implantation of human MC-Chs and NCC-Chs in mice. **(A)** Depiction of the experimental procedure of *in vitro* chondrogenic induction and subcutaneous implantation of MC-Ch and NCC-Ch pellets. Macrographs and size quantification of cell pellets before and after implantation were shown. **(B)** Implanted MC-Ch and NCC-Ch pellets analyzed by H&E and Safranin O staining. Dash-boxed areas in the left column are shown at a higher magnification in the right column for each staining. **(C)** Immunofluorescence staining of implanted MC-Ch and NCC-Ch pellets for detection of human vimentin to distinguish implanted human cells from host mice cells. Nuclear DNA was labeled with DAPI. **(D)** Immunofluorescence staining of implanted MC-Ch and NCC-Ch pellets for detection of COL2A1 and COL10A1. Nuclear DNA was labeled with DAPI. *n* = 3. Scale bar, 200 μm.

### MC-Ch and NCC-Ch pellets repair joint defects of rats

To further examine the capacity of cartilage generation, pellets of MC-Chs and NCC-Chs were implanted in osteochondral defects and sealed with fibrin glue at the femoral trochlea groove of athymic nude rats (Fig. 3A). The MC-Ch and NCC-Ch groups were then compared to the negative control group of fibrin glue, in which a defect was sealed without pellet implantation. Macroscopic evaluation showed that defects of both the MC-Ch and NCC-Ch groups were filled with regenerated tissue after 4 weeks, whereas those of the fibrin glue group were not (Fig. 3B). At week 16, increased joint repair was found in all groups with the NCC-Ch group exhibiting more complete repair than the other groups.

**Fig. 3.**
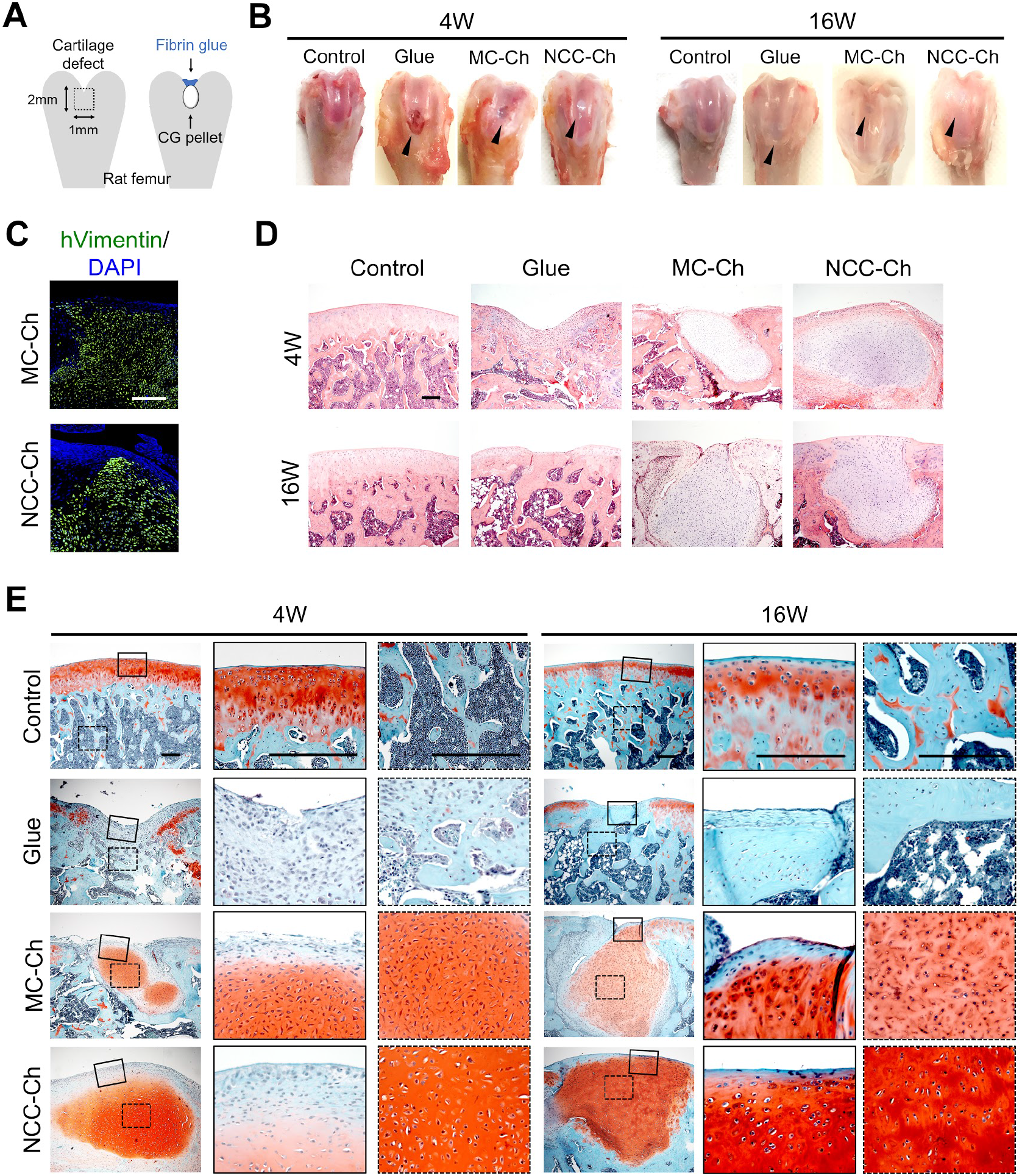
Implantation of human MC-Ch and NCC-Ch pellets in rat joints. **(A)** Schematic of chondrocyte pellet implantation. The pellet placed in the joint defect was covered with fibrin glue for secure integration with host tissue. **(B)** Repair of cartilage defects 4 weeks and 16 weeks after pellet implantation. The control shown is contralateral joints of animals receiving implants. Black arrows point to the site of implantation. **(C)** Immunofluorescence staining of MC-Ch and NCC-Ch pellets for detection of human vimentin. Nuclear DNA was labeled with DAPI. **(D)** Rat femur joints analyzed by H&E staining. **(E)** Rat joints analyzed by Safranin O staining. Solid-boxed (superficial) and dash-boxed (subchondral bone) areas in the left column are shown at a higher magnification in the central and right column, respectively. *n* = 3. Scale bar, 200 μm.

Human vimentin localization indicates that the repair tissue at joint defects of rats implanted with MC-Ch and NCC-Ch pellets comprised mainly human cells (Fig. 3C). The H&E staining results showed that in the fibrin glue group, fibrous tissue resulting from host cells was formed after 4 weeks and later replaced by bone-like tissue after 16 weeks (Fig. 3D). On the other hand, the groups implanted with cell pellets showed that the defects were filled with highly cellular tissue after 4 and 16 weeks. Particularly, the regenerated tissue derived from NCC-Ch pellets integrated well with the surrounding native cartilage at week 16 (Fig. 3D). Safranin O staining further revealed that the regenerated tissue by MC-Chs or NCC-Chs was rich in GAG content and contained lacunae with spindle-shaped or round-shaped cells, respectively, but both groups fell short of fully restoring the osteochondral structure of cartilage as that shown in the control (Fig. 3E). These results demonstrate the capacity of MC-Chs and NCC-Chs for cartilage regeneration of joint defects and a more desired outcome of cartilage repair by the NCC-Ch group.

### Global transcriptome of MC-Chs and NCC-Chs largely resembles that of NCs

To further investigate cell differences at the molecular level, the global transcriptome of MC-Chs and NCC-Chs, compared to that of NCs, was analyzed by RNA sequencing. Overall differences in the transcriptome between MC-Chs or NCC-Chs and NCs were revealed by the Euclidean distance in multidimensional scaling (MDS) plots (Fig. 4A) and also demonstrated by hierarchical clustering heatmaps (Fig. 4B). About 8% or 6% of the total transcript was differentially expressed between MC-Chs or NCC-Chs and NCs, respectively, and 5% of the total transcript between MC-Chs and NCC-Chs (Fig. 4C and fig. S3A). The differentially expressed transcripts with the information of fold change, *p* value, and false discovery rate (FDR) are listed in Tables S1-S3. The Gene Ontology (GO) enrichment result showed that transcripts associated with lineage-specifications, including epithelium development and skeletal system development, were differentially regulated in MC-Chs, whereas those, including neurogenesis and nervous system development, were up-regulated in NCC-Chs compared to NCs (Fig. 4D). A number of transcripts highly expressed in NCC-Chs compared to those in MC-Chs were neuron development-related (fig. S3B). Further analyzed using the KEGG pathway database, the result showed that the transcripts up-regulated in both MC-Chs and NCC-Chs, compared to NCs, were related to complement and coagulation cascades and glutathione metabolism (Fig. 4E), previously found involved in the regulation of chondrogenesis (*22, 23*). The transcripts of molecules in the pathways mediating tight junction and focal adhesion in MC-Chs and cell adhesion molecules and ECM receptor interaction in NCC-Chs, critical to cell condensation of chondrogenesis, were up-regulated compared to those in NCs. In addition, the result showed that transcripts of molecules in the pathways controlling cholesterol metabolism, PPAR signaling pathway, and fat digestion and absorption in MC-Chs were increasingly expressed compared to those in NCC-Chs (fig. S3C), which were previously found related to the phenotype of growth plate chondrocytes (*24–26*). Altogether the bioinformatic results indicate that both MC-Chs and NCC-Chs are phenotypically similar but not identical to NCs.

**Fig. 4.**
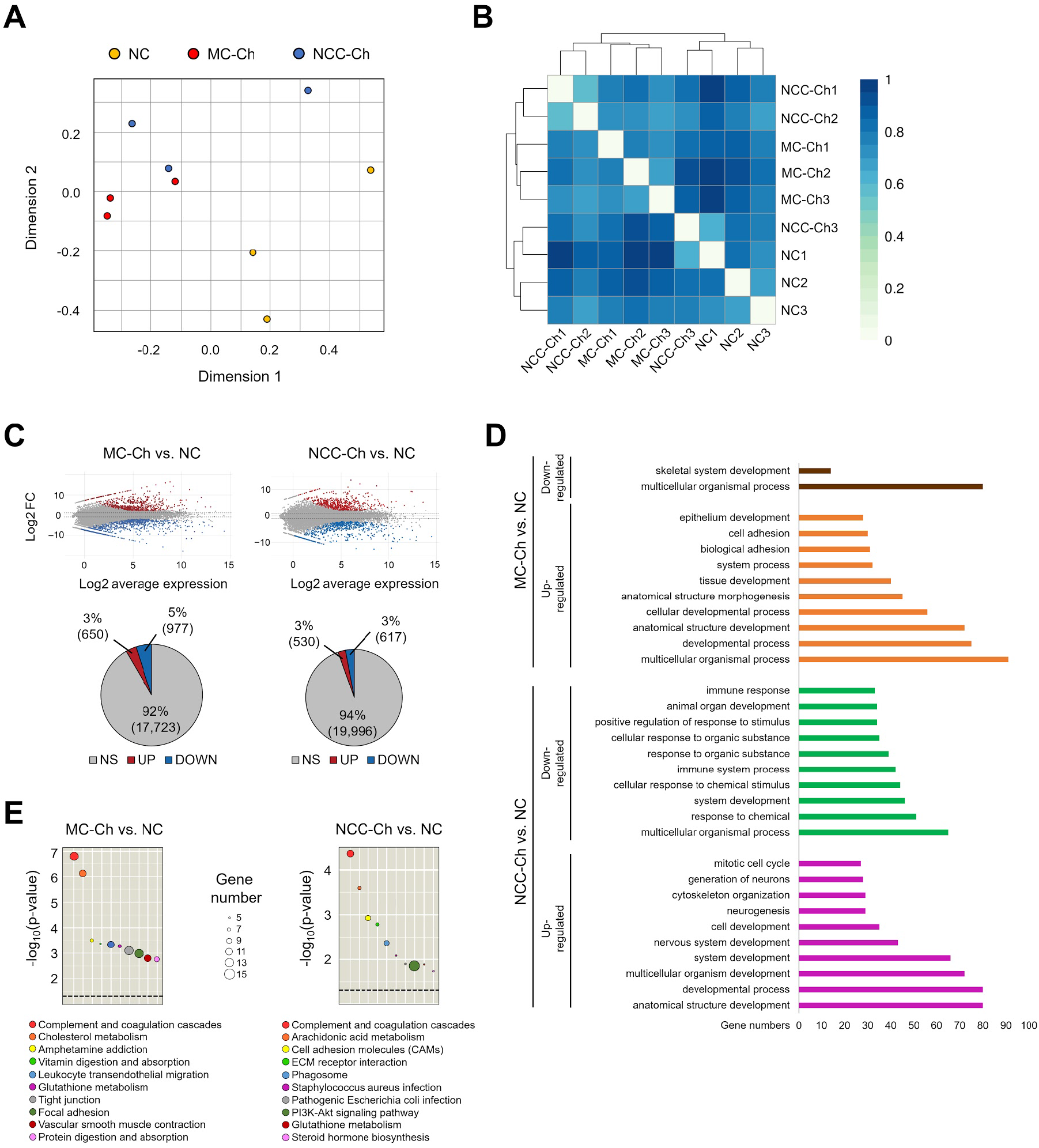
Comparison of global transcriptome profiles between hiPSC-derived and native chondrocytes. **(A)** MDS plot depicting visualization of principal component analysis and variations in MC-Chs, NCC-Chs and NCs. **(B)** Relationships between each sample group shown in a heatmap. Each entry is colored on the basis of its dissimilarity to other samples, and the rows and columns are reordered according to the hierarchical clustering. A dendrogram depicts the hierarchical arrangement of the samples produced by the hierarchical clustering. *n* = 3. **(C)** Smearplots presenting differences in the global transcript expression of MC-Chs or NCC-Chs compared to NCs. Red and blue dots denote individual up- and down-regulated genes, respectively, at the significance threshold of adjusted *p*-value of 0.05. Gray dots reflect those genes with no statistically significant differential expression. Percentages of the up-regulated (Up), down-regulated (Down), and non-significant (NS) genes are shown in corresponding pie charts. **(D)** Gene ontology enrichment analysis of MC-Chs or NCC-Chs compared to NCs. Categories of the most differentially expressed gene sets with a *p*-value less than 0.05 are shown. The number of genes in each enrichment category is represented by the length of an individual bar. **(E)** Analysis of KEGG pathway enrichment identifying abundance of differentially expressed genes in predominant biological categories of MC-Chs or NCC-Chs compared to NCs. Significantly enriched KEGG pathways are ordered from most to least significant. The number of genes in a specified KEGG pathway is denoted by the size of a circle. The color of each circle corresponds to that of the specific KEGG pathway. The dashed line represents significance at *p*-value = 0.05. *n* = 3.

### Transcripts of cartilage development-associated molecules are differentially expressed between MC-Chs or NCC-Chs and NCs

Cartilage development-associated genes (GO: 0051216) of the GO database were selected to further characterize the differentially expressed transcripts between MC-Chs, NCC-Chs, and NCs. The analysis showed that 34% and 29% of the cartilage development-associated transcripts were significantly regulated in MC-Chs and NCC-Chs, respectively, compared to those in NCs (Fig. 5A and 5B) while 11% of the transcripts between MC-Chs and NCC-Chs were differentially expressed (fig. S4A). Heatmaps of differential expression of the transcripts between MC-Chs, NCC-Chs, and NCs showed that many of the listed molecules are associated with chondrogenesis and ECM organization (Fig. 5C, 5D, and fig. S4B). Analysis of functional protein association networks revealing interactions of the differentially expressed transcripts determined that transcription factors and growth factors were the most interconnected molecules in the networks (fig. S4C and fig. S5A, 5B). In particular, BMP2, BMP6, and FGF2 were highly interactive in the networks of 54 transcripts downregulated in MC-Chs, and BMP2, BMP6, FGF2, and GDF5 in the networks of 45 transcripts downregulated in NCC-Chs, compared to those in NCs (fig. S5A and S5B). These results indicate that chondrocytes derived from hiPSCs using the induction protocols described in Figure 1A are close but not identical to phenotypic articular chondrocytes, suggesting that the protocols require further improvement to reduce the phenotypic differences between MC-Chs or NCC-Chs and NCs.

**Fig. 5.**
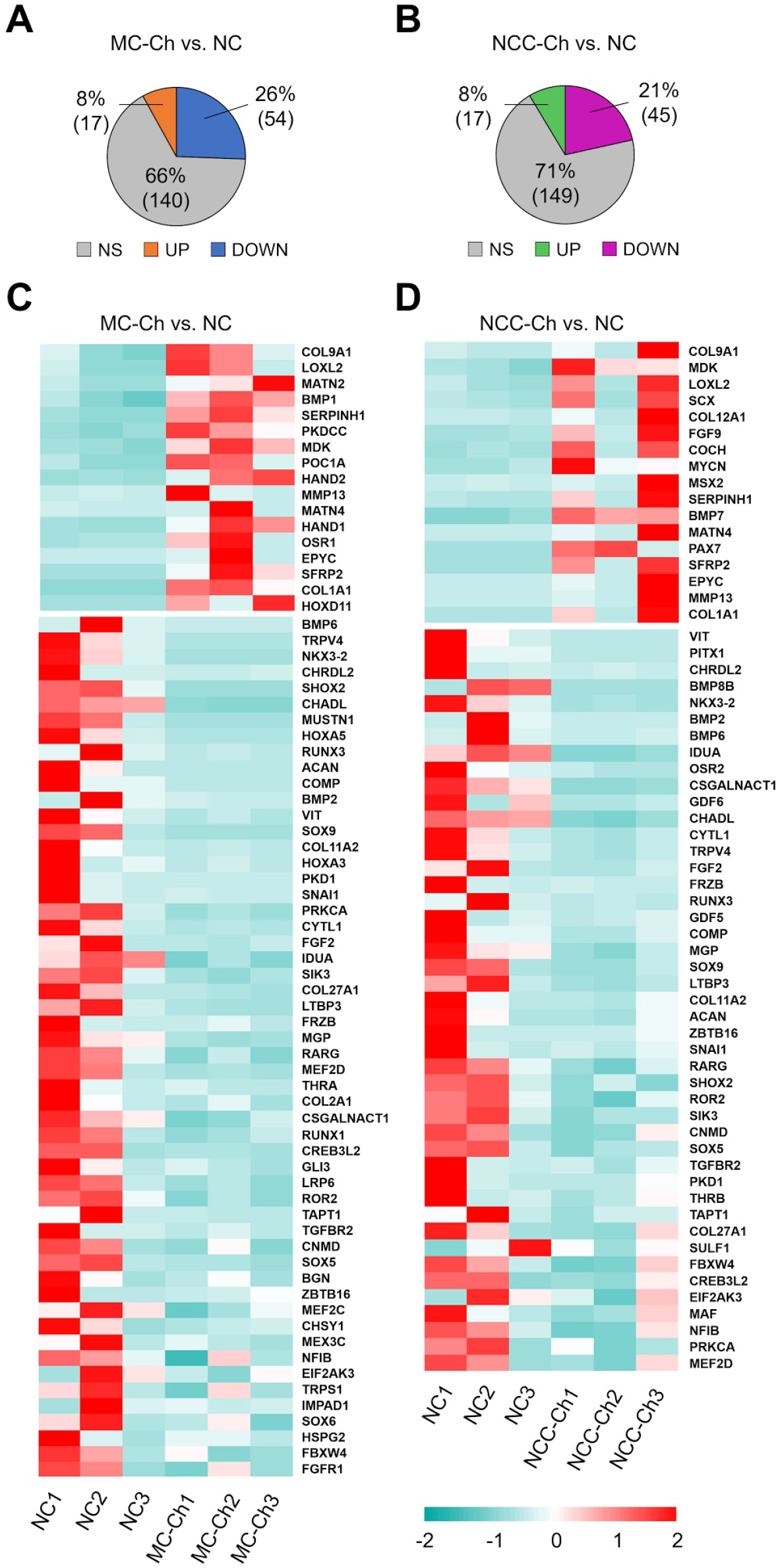
Analysis of differentially expressed transcripts associated with cartilage development between hiPSC-derived and native chondrocytes. **(A-B)** Pie charts depicting percentages of the up-regulated (Up), down-regulated (Down), and non-significant (NS) transcripts in MC-Chs or NCC-Chs compared to NCs. **(C-D)** Heatmaps of differentially expressed cartilage development-associated transcripts between MC-Chs or NCC-Chs and NCs. The color intensity of each grid denotes the extent of change in transcript expression, where up-regulated transcripts are shown in red and down-regulated ones in turquoise. *n* = 3.

### Chondrogenic induction with selected growth factors enhances differentiation of hyaline chondrocytes from MCs and NCCs

To test the hypothesis that the growth factors, downregulated in MC-Chs or NCC-Chs compared to NCs (fig. S5), are essential to induction of phenotypic articular chondrocytes, we induced chondrogenic differentiation of MCs and NCCs using TGFB1-based medium with or without a combination of the selected growth factors (Fig. 6A). After 21 days of chondrogenic differentiation, MC-Ch and NCC-Ch pellets of the treated groups exhibited a significant increase in size (Fig. 6B), GAG content (Fig. 6C), and intensity of Alcian blue staining (Fig. 6D) compared to those of the control groups. The result of mRNA expression showed that hyaline cartilage-associated markers, *SOX9*, *ACAN*, and *COL2A1*, were upregulated during chondrogenic induction of MCs and NCCs and exhibited higher levels in treated MC-Chs and NCC-Chs than those in control cells (Fig. 6E). Notably, expression levels of articular cartilage-associated markers, *SIX1, THBS4*, and *ABI3BP (27),* were significantly higher in treated MC-Chs than those in control cells whereas treated NCC-Chs did not show the similar response. These results seem to indicate that NCC-Chs induced by a combination of the growth factors may become hyaline craniofacial chondrocytes other than articular ones. Interestingly, the expression of *COL1A1,* a marker of fibrocartilage, was significantly reduced in NCC-Chs in response to the treatment whereas that in MC-Chs remained comparable between the control and treated groups. Markers of hypertrophic chondrocytes, *COL10A1* and *ALPL,* were downregulated in treated MC-Chs compared to those in control cells; in contrast, expression levels of these two markers in treated NCC-Chs were higher than those in the control. Results of immunofluorescence staining revealed that the COL2A1 content in treated MC-Ch and NCC-Ch pellets was comparable to that in control ones (Fig. 6F). Consistent with the mRNA expression results, COL10A1 and COL1A1 were significantly reduced in treated MC-Ch and NCC-Ch pellets, respectively, compared to that of control ones. It is worth noting that the increase in the expression of *COL10A1* transcripts between treated and control NCC-Chs was not shown in the protein expression. Taken together, these results demonstrate that our modified protocols using the growth factors identified in this study promote chondrogenesis of MCs and NCCs toward the generation of hyaline cartilage chondrocytes.

**Fig. 6.**
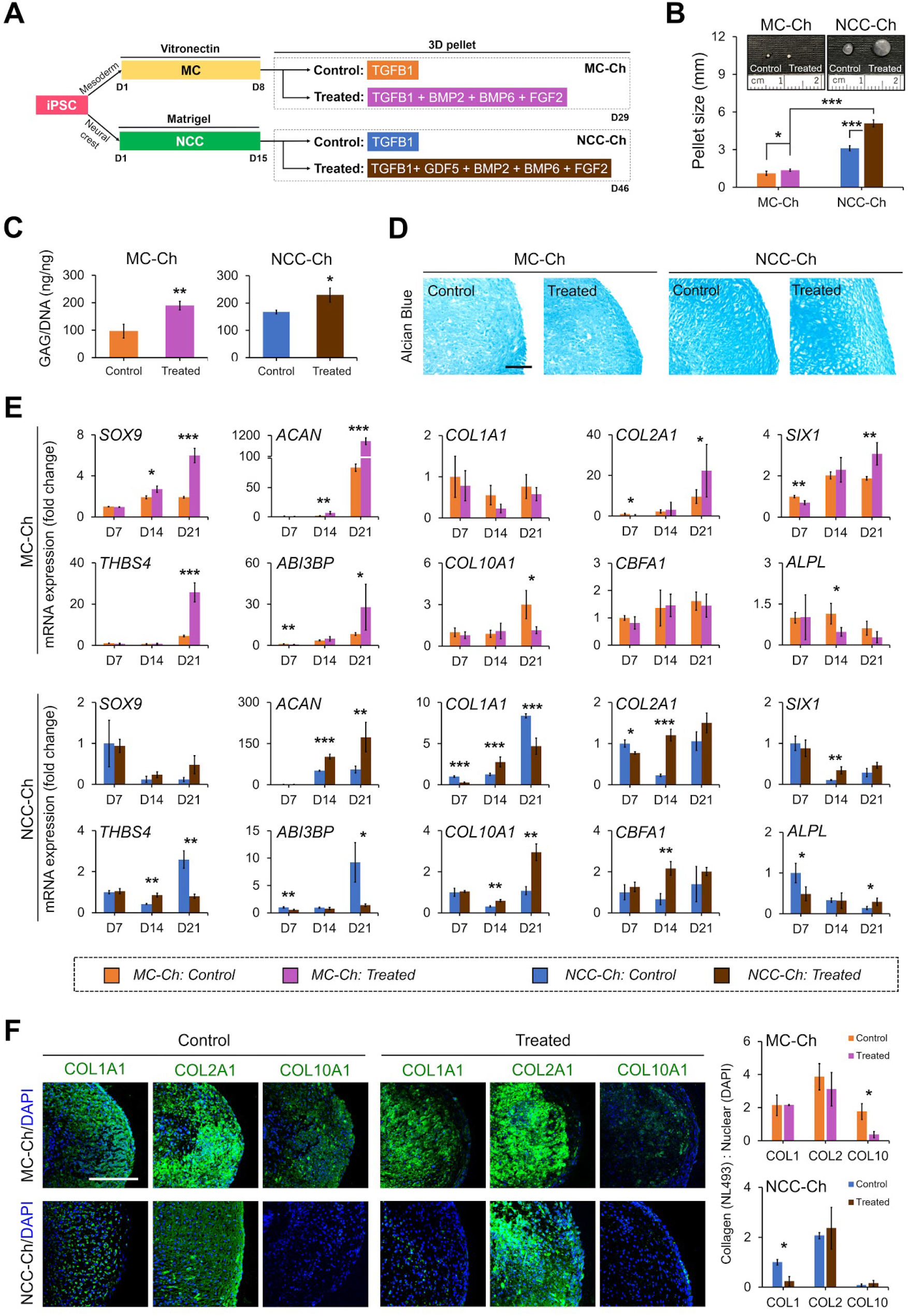
Chondrogenic induction of MCs and NCCs by newly identified growth factors. **(A)** Schematic of procedures inducing differentiation from hiPSCs to chondrocytes and corresponding timelines. **(B)** Macrographs and size quantification of representative MC-Ch and NCC-Ch pellets of control and treated groups. **(C)** GAG quantification of MC-Ch and NCC-Ch pellets of control and treated groups. **(D)** Alcian blue staining of MC-Ch and NCC-Ch pellets of control and treated groups. **(E)** Levels of mRNA expression of general cartilage-associated markers (*SOX9*, *ACAN, COL1A1, COL2A1*), hyaline cartilage-associated markers (*SIX1, THBS4, ABI3BP*) and hypertrophic-associated markers (*COL10A1, CBFA1, ALPL*) in MC-Ch and NCC-Ch pellets. **(F)** Immunofluorescence staining of MC-Ch and NCC-Ch pellets of control and treated groups for detection of COL1A1, COL2A1, and COL10A1. Nuclear DNA was labeled with DAPI. *n* = 3. *\*p* < 0.05, *\*\*p* < 0.01, *\*\*\*p* < 0.001. Scale bar, 200 μm.

## Discussion

Research focusing on derivation of chondrocytes from iPSCs for cartilage regeneration has gained increasing attention in recent years, though it still remains challenging to obtain phenotypic articular chondrocytes (*28*). In this study, the phenotype, chondrogenic capability, and transcriptome of isogenic MC-Chs or NCC-Chs differentiated from blood-derived hiPSCs along the mesodermal and ectomesodermal lineage, respectively, were characterized. The *in vitro* results have shown that NCC-Chs express greater levels of hyaline cartilage-associated markers and produce more GAG production compared to MC-Chs. Notably, NCC-Chs do not express hypertrophic chondrocyte-associated markers. Results of *in vivo* studies have further shown that the articular cartilage layer with articular chondrocyte-like cells and proteoglycan-rich ECM is regenerated from implanted NCC-Chs at a joint defect. To the best of our knowledge, the direct comparison between isogenic chondrocytes derived from the mesodermal and neural crest lineages has not been reported. Moreover, we have demonstrated that the derivation of both MC-Chs and NCC-Chs is enhanced by a combination of cartilage development-associated growth factors identified in this study.

We derive MC-Chs and NCC-Chs from hiPSCs using previously published protocols with our modification. While our findings are largely in agreement with those reported by other groups, there exist differences between their and our methods and results. To derive chondrocytes along the mesodermal lineage, we follow the protocol published by Kimber’s group (*9*) to induce hiPSCs into MCs. Instead of culturing MCs on a gelatin-coated two-dimensional (2D) surface with induction of FGF2 and GDF5 in their study, we pack the cell in a three-dimensional (3D) cell pellet to increase cell-cell interaction and induce chondrogenesis with TGFB1. The modified protocol is adopted to address concerns associated with the use of a 2D surface for chondrogenesis. A previous study has shown that chondrocytes generated from hiPSCs in 2D culture contain undifferentiated hiPSCs and implantation of the chondrocyte at a joint defect results in teratoma formation (*12*). The findings indicate that complete chondrogenesis is not achieved in 2D culture due to lack of cell condensation, critical for early chondrogenic induction and cartilage formation (*29*). In our current study, generated MC-Chs synthesize noticeable COL1 and COL10 in pellets (Figs. 1H and 2D) whereas the cell derived by Kimber’s group expresses negligible mRNA levels of the molecules (*15*). The difference likely resulted from the effect of FGF2, given that our current results show that the modified induction medium containing FGF2 significantly reduces COL10 in a chondrogenic pellet (Figs. 6E and F), supported by a previous finding that FGF2 is able to ameliorate chondrocyte hypertrophy (*30*). For derivation of NCC-Chs, we follow our previously reported protocol (*20*) to induce differentiation of hiPSCs along the ectomesodermal lineage. Chondrocytes derived from NCCs express abundant COL2 and GAG but little COL1 and no COL10 (Figs. 1H and 2D). The current finding is discrepant with previously reported results that hESC-derived NCCs induced by PDGF and BMP4 increasingly express COL10 during chondrogenesis and subcutaneous implantation of the cell leads to bone formation (*31*). Altogether, these results suggest that it is critical to optimize the culture condition for inducing chondrogenesis without leading to chondrocyte hypertrophy.

One of the key findings in this study is that 92% and 94% of the mRNA transcript of MC-Chs and NCC-Chs, respectively, show expression levels comparable to that of NCs, suggesting that the hiPSC-derived cells are similar but not identical to phenotypic articular chondrocytes. Specifically, our results indicate that the differentially expressed genes between MC-Chs and NCs are associated with epithelium development and blood vessel formation, whereas those between NCC-Chs and NCs are associated with neuron generation and nervous system development. The similar finding that stem cell-derived chondrocyte culture contains multiple cell populations has been reported by other groups. For example, using the approach of single cell analysis, Dicks *et al*. have found that at least 9 different cell clusters, including chondrogenic, neurogenic, other non-bone/non-cartilage mesenchymal lineage cells, are identified in hiPSC-derived mesodermal chondrocyte culture (*32*). Similarly, Umeda *et al*. have demonstrated that the culture of ectomesodermal chondrocytes derived from hESCs and hiPSCs contains cell populations capable of differentiating into neural cells, melanocytes, and endothelial cells (*18*). While it has remained challenging to derive a homogenous population of target cells from PSCs (*33, 34*), our approach to identifying additional induction molecules to enhance chondrogenesis offers a potential solution.

The analysis of differential expression of cartilage development-associated genes reveals differences between hiPSC-derived and native chondrocytes, indicating that TGFB alone is inadequate to induce complete chondrogenesis of hyaline cartilage. TGFB1 and TGFB3 have been extensively used as potent growth factors for chondrogenic induction of stem cells, but they have been shown to increase endochondral ossification by upregulating *RUNX2* expression (*35*). In addition, studies have reported that TGFBs play a role in the formation of fibrocartilage (*36, 37*). These findings are consistent with our results shown in Figures 1H and 2D. In this study, we have demonstrated that with the supplementation of BMP2, BMP6, FGF2, and GDF5 in TGFB1-based induction medium, the phenotype of MC-Chs and NCC-Chs is increasingly driven toward that of hyaline cartilage chondrocytes.

We have found that FGF2 and BMPs play a critical role in the generation of hyaline cartilage. Our results show that the size of cell pellets treated with TGFB1 and the added growth factors is markedly larger than that with TGFB1 alone. The increase in pellet size is likely resulted from the effect of FGF2 on chondrogenesis. We and others have shown that FGF2 promotes cell proliferation and increases *COL2* expression in chondrogenic cells (*38, 39*). Previous studies have also shown that BMPs are involved in chondrogenesis and the development of hyaline cartilage. For example, BMP2 enhances COL2 and aggrecan production in chondrocytes (*19*), BMP6 promotes chondrogenic induction and hyaline cartilage regeneration (*40*), and GDF5, also known as BMP14, enhances the expression of *SOX9*, *COL2,* and *ACAN* and significantly reduces the expression of hypertrophic chondrocyte-associated markers (*41*). In addition to these effects, BMPs have been shown to reduce fibrous tissue formation. For instance, BMP2 is able to ameliorate TGFB-induced fibrosis in liver (*42*), and BMP6 downregulates COL1 expression to reduce fibrocartilage formation (*40*). These findings may explain our result that the addition of BMPs reduces hypertrophy of MC-Chs and fibrosis of NCC-Chs. Nevertheless, our results indicate that further optimization of the induction medium may be required for enhancing phenotypes of hiPSC-derived chondrocytes. Specifically, Figures 6E and F show that there is discrepancy between the mRNA and protein results of hypertrophic chondrocyte-associated markers of NCC-Chs treated with and without the combined growth factors. Although the transcript result alone is inadequate to confirm hypertrophy of NCC-Chs, it still indicates the cell may be over-induced by non-optimal doses of the growth factors to undergo the early stage of hypertrophy. To overcome the issue, our future study will focus on optimizing induction doses and timing of the growth factors during chondrogenesis of NCCs and MCs to generate phenotypic hyaline-cartilage chondrocytes.

Our current study uses articular chondrocytes as the reference cell to determine differences in the expression of transcriptome between hiPSC-derived and target chondrocytes. We are aware that our approach to compare ectomesodermal NCC-Chs and mesodermal articular chondrocytes may be less-then-idea for finding developmentally relevant molecules to enhance NCC chondrogenesis. However, considering that our purpose is to generate articular cartilage and it would be challenging to correctly compare MC-Chs and NCC-Chs with different reference cells, we believe that our experimental design using the same type of reference chondrocytes is properly justified. It would be of interest for future studies to use native chondrocytes isolated from craniofacial cartilage as reference cells in comparison with ectomesodermal lineage-derived chondrocytes. This may allow identification of different sets of growth factors specific for induction of craniofacial chondrocytes of hyaline cartilage.

The main finding is that using the protocols described in this study, chondrocytes derived from the ectomesodermal lineage share more cellular and molecular characteristics of hyaline cartilage chondrocytes and have a greater capability to generate cartilage compared to those derived from the mesodermal lineage. Indeed, the use of craniofacial chondrocytes for joint repair has been demonstrated by several groups. Studies have shown that transplanted autologous nasal chondrocytes are able to repair articular cartilage of large animals and successfully reconstruct joint surfaces in a first-in-human trial (*43, 44*). Similarly, another study has demonstrated that an osteochondral defect of a rabbit is repaired by autologous nasal chondrocytes with neo-tissue highly similar to articular cartilage (*45*). Together, these studies provide compelling evidence to support the potential application of ectomesodermal lineage-derived cells for cartilage repair.

## Materials and Methods

### Culture of blood-derived hiPSCs

Three independent blood-derived hiPSC lines, JHU-62i, JHU-106i, and JHU-160i, derived from different donors were obtained from the WiCell Laboratory (Madison, WI, USA). These cell lines were cultured with Essential 8 (E8) medium in vitronectin-coated 6-well plates, and maintained in an incubator at 37°C in a humidified 5% CO_2_ atmosphere. Cells were stained with Fast blue RR salt (Sigma-Aldrich, St. Louis, MO, USA) to detect localization of ALP. When reached about 80% confluence, cells were collected using Versene solution (Thermo Fisher Scientific, MA, USA) and re-plated at a split ratio of 1:6. hiPSC lines at Passage 12 were used in this study.

### Generation of human iPSC-derived MCs and NCCs

To obtain distinct lineage-derived chondrocytes, individual hiPSC lines were induced into mesodermal and ectomesodermal lineages following previously published protocols by Ordershaw *et al*. (*9*) and our collaborative group (*20*), respectively. For MC induction, hiPSCs were seeded in fibronectin (Thermo Fisher Scientific) coated 6-well plates and induced by mesodermal differentiation basal medium (DMEM/F12, 2mM of L-glutamine, 1% non-essential amino acid, 1% ITS, 2% B27, and 90 uM of beta-mercaptoethanol) composed of a combination of chemicals with different dosages, as listed in Table S4, dependent on induction stages. Briefly, in the initial stage of 4 days, hiPSCs were induced into mesendodermal cells by basal medium containing 10-50 ng/ml of Activin-A, 25 ng/ml of WNT3A, 20 ng/ml of FGF2, and 40 ng/ml of BMP4. In the next 4 days, mesendodermal cells were then induced into MCs with basal medium containing 20 ng/ml of FGF2, 40 ng/ml of BMP4, 100 ng/ml of Follistatin, and 2 ng/ml of NT4. On the other hand, ectomesodermal NCCs were differentiated from hiPSCs by simultaneously inhibiting SMAD and activating WNT signaling (*20*). Briefly, hiPSCs seeded at a density of 67,500 cells/cm2 were induced by differentiation medium containing Essential 6 (E6) medium, 5 mM of SB431542, 10 mM of dorsomorphin, 30 mg/ml of heparin, 10 mM of CHIR99201, and 100 ug/ml of FGF2 for 12 days. Cells were passaged using Accutase (Stemcell Technologies, Vancouver, BC, Canada) when reaching about 90% confluence. To prepare NCCs, passaged cells were washed 3 times with ice-cold Dulbecco’s Phosphate Buffered Saline (D-PBS) washing buffer, treated with Neural Crest Stem Cell MicroBeads for 30 min at 4°C, and sorted by magnetic-activated cell sorting. The sorted NCCs were then seeded and expanded in growth factor-reduced Matrigel (WiCell Laboratory)-coated 6-well plates for subsequent chondrogenic induction.

### *In vitro* chondrogenic induction and assessment

To prepare mesoderm- and ectomesoderm-derived chondrocytes, MCs and NCCs were further induced for chondrogenesis following our previously published protocol (*46*). Briefly, 500,000 MCs or NCCs were collected and centrifuged at 600 *g* for 5 min to form a high-cell-density pellet before induced by chondrogenic medium containing high-glucose DMEM, 1% antibiotics, 1% ITS+ Premix (Corning), 0.9 mM of sodium pyruvate, 50 μg/ml of L-ascorbic acid-2-phosphate, 40 μg/ml of L-proline, 0.1 μM of dexamethasone, and 10 ng/ml of TGFB1 (PeproTech, Rocky Hill, NJ, USA). Medium was changed every 2 days during the period of chondrogenic induction. To further enhance the derivation of phenotypic chondrocytes, MC and NCC pellets were induced by TGFB1-based differentiation medium with 10 ng/ml of BMP2, BMP6, and FGF2 each. For chondrogenesis of NCC pellets, 10 ng/ml of GDF5 was additionally added. Chondrogenesis of cells by TGFB1 alone was used as a control.

For analysis of chondrogenic differentiation, chondrocyte pellets were fixed in 4% formaldehyde, dehydrated by a series of gradient ethanol, infiltrated with xylene, and then embedded in paraffin. Embedded cell pellets were cut into 8-μm sections using a microtome, deparaffinized, and stained with Alcian blue (Polysciences,Warrington, PA, USA) for GAG detection. In addition, the total GAG content of cell pellets was quantified by the dimethylmethylene blue assay (Sigma-Aldrich) and normalized with the DNA content determined separately by the PicoGreen assay (Thermo Fisher Scientific) following the manufacturer’s instructions.

### Native chondrocytes harvested from human articular cartilage

Human articular cartilage as part of surgical waste obtained from 3 donors without osteoarthritis was provided by the University of Wisconsin Hospital and Clinics with approval of the Institutional Review Board. To isolate chondrocytes, cartilage was individually minced, washed twice with D-PBS, and incubated with 1 mg/ml of pronase solution (Roche, Basel, Switzerland) for 30 min at 37°C and additionally with 1 mg/ml of collagenase solution (Roche) for 12-18 hr at 37°C with agitation. The digestion solution was collected and centrifuged at 600 g for 5 min to prepare uncultured native chondrocytes for RNA-seq analysis.

### Flow cytometry analysis

Cells were suspended and washed 3 times with washing buffer composed of ice-cold D-PBS, 1% BSA, 5mM of EDTA, and 25mM of HEPES before antibody incubation. hiPSCs were incubated with the primary antibody goat anti-human OCT3/4 (R&D Systems, Minneapolis, MN, USA), mouse anti-human SOX2 (R&D Systems), goat anti-human NANOG (R&D Systems), mouse anti-human PODPCALYXIN (R&D Systems), mouse anti-human SSEA4 (R&D Systems), or mouse anti-human CD9 (R&D Systems) for 30 min at 4°C. After three subsequent washes with the buffer, cells were incubated with secondary antibody Alexa Fluor-546 donkey anti-mouse (Thermo Fisher Scientific) or FITC donkey anti-goat (Santa Cruz Biotechnology, Santa Cruz, CA) for another 30 min at 4°C. For analysis of MC-Chs and NCC-Chs, cells were incubated with PE-conjugated mouse anti-human CD44 or CD151 (BD) for 30 min at 4°C. Fluorescent cells were then detected by the Attune™ NxT Flow Cytometer (Life Technologies, Carlsbad, CA), and the resulting flow cytometry data were analyzed by FlowJo (Treestar, Ashland, OR), following the manufacturer’s instructions.

### Immunofluorescence analysis

hiPSCs were fixed with 4% formaldehyde for 20 min at room temperature, and permeabilized with 0.1% Triton X-100 in D-PBS for 5 min at 4°C before incubated with the primary and secondary antibodies previously described in the method of flow cytometry analysis. Cell pellets harvested from chondrogenic culture or implants from animals were cut into 8-μm sections, deparaffinized, and unmasked with 0.1% (wt/vol) pepsin in 0.01N HCl for 15 min, permeabilized with 0.1% Triton X-100 in D-PBS for 30 min, and blocked with 0.1% (wt/vol) BSA in D-PBS for 30 min. Slides with sectioned specimens were incubated with primary antibody goat anti-human COL1A1 (Santa Cruz Biotechnology), COL2A1 (Santa Cruz Biotechnology), COL10A1 (Santa Cruz Biotechnology), or rabbit anti-human vimentin (Abcam, Cambridge, UK) for an hour at 4°C. The slides were then washed three times with ice-cold D-PBS washing buffer, incubated with secondary antibody FITC donkey anti-goat (Santa Cruz Biotechnology) or Alexa Fluor-555 donkey anti-rabbit (Thermo Fisher Scientific) for 45 min at 4°C, and mounted with coverslips using mounting medium with DAPI (Vector, Burlingame, CA). The dilution of antibodies for optimal concentration was set following individual manufacturer’s instructions. For fluorescence detection, specimens were observed under the Nikon A1R-s Confocal Microscope (Nikon, Tokyo, Japan).

### Total RNA extraction and real-time qPCR

Total RNA was extracted from cells using the NucleoSpin RNA II kit (Clontech Laboratories, Mountain View, CA, USA). The quality and quantity of total RNA were measured by Nanodrop 1000 (Thermo Fisher Scientific), followed by reverse transcription with the High-Capacity cDNA Reverse Transcription kit for real-time qPCR. To quantify the mRNA expression of markers of interest, cDNA was amplified by qPCR with iQSYBR Green Premix (Bio-Red, Hercules, CA, USA) and specific primers listed in Table S5. The level of mRNA expression was determined using the 2^-Δ*Ct*^ method with *ubiquitin C* as a housekeeping gene.

### *In vivo* evaluation of human iPSCs and derivatives

*In vivo* studies were performed following the animal protocols approved by the University of Wisconsin-Madison Institutional Animal Care and Use Committee. For the teratoma formation assay, 1 million hiPSCs in 0.1 ml Matrigel were mixed with DMEM/F12 at the ratio of 1:1, and subcutaneously injected into the hind leg of SCID mice (NOD.Cg-Prkdc^scid^I12rg^tm1wj1^/SzJ) (The Jackson Laboratory, Bar Harbor, ME, USA). Teratoma tissue was harvested eight weeks after injection and prepared for histological analysis by H&E staining. The same strain of mice was also used for ectopic implantation of MC-Ch and NCC-Ch pellets. Briefly, a 10 mm incision was created for insertion of a single pellet into the hind leg of a mouse. After 30 days of subcutaneous implantation, pellets were collected and processed for immunohistochemical and H&E and Safranin O staining.

To evaluate the capacity of hiPSC-derived chondrocytes for articular cartilage repair, MC-Ch and NCC-Ch pellets were implanted into joints of 18 athymic nude rats (Hsd:RH-Foxn1^mu^) that were equally divided into 4- and 16-week treatment groups with each further divided into acellular fibrin glue (n = 3), MC-Ch pellet (n = 3), and NCC-Ch pellet (n = 3) subgroups. A cylindrical defect of 1 mm in diameter and 2 mm in depth was created by a biopsy punch at the right femur groove of a rat for implantation, and the left joint of the animal was used as a contralateral control. Tissue specimens repaired by cell pellets were harvested after animals were euthanized, fixed with 4% formaldehyde, and decalcified with 22% formic acid prior to paraffin embedding and sectioning for the analysis of immunohistochemical and histological staining and microscopic analysis (Nikon, Japan).

### RNA-sequencing assessment and data analysis

Native chondrocytes, MC-Chs, and NCC-Chs were collected, lysed in Trizol (Life technologies), and stored in −80°C until extraction of total RNA extraction using the Direct-zol^™^ RNA kit (Zymo Research). The optical density value of total RNA was measured using the NanoDrop spectrophotometer (Thermo Fisher Scientific) to confirm an A260:A280 ratio above 1.9, and the RNA integration number (RIN) was measured using the BioAnalyzer (Agilent Technologies) RNA 6000 Nano kit. Complementary DNA libraries were prepared using the SMART-Seq v4 Ultra Low Input RNA kit for Sequencing (TAKARA Bio) and Nextera XT DNA Library Prep kit (Illumina). Size distribution and concentration of the final products were then evaluated using the Bioanalyzer High Sensitivity DNA kit (Agilent Technologies). The libraries were pooled and diluted to 3 nM using 10 mM Tris-HCl, pH 8.5 and then denatured using the Illumina protocol. The denatured libraries were loaded onto an S1 flow cell on Illumina NovaSeq 6000 (Illumina) and ran for 2×50 cycles according to the manufacturer’s instructions. De-multiplexed sequencing reads were generated using Illumina bcl2fastq (released version 2.18.0.12), allowing no mismatches in the index read.

BBDuk (REF) was used to trim/filter low quality sequences using the “qtrim=lr trimq=10 maq=10” option. Next, alignment of the filtered reads to the human reference genome (GRCh38) was done using HISAT2 (version 2.1.0) (REF) applying “--no-mixed and --no-discordant’’ options. Read counts were calculated using HTSeq (REF) by supplementing Ensembl gene annotation (GRCh38.78). Gene expression values were calculated as counts per million (CPM). Genes with no detected CPM in all samples were filtered out.

To identify the transcripts and associated expression levels, the genomic origin of sequenced cDNA fragments must be determined. Raw fastq files were preprocessed with the trimming software Skewer (*47*), and the trimmed single-end reads were aligned to the reference human genome using STAR (Spliced Transcripts Alignment to a Reference) (*48*). In addition, samples were normalized by the method of trimmed mean of M-values (TMM) (*49*), and the analysis of differentially expressed genes was performed with a generalized linear model using the edgeR package (*50*). All genes with a *p*-value < 0.05 and FDR < 0.25 were selected to conduct the test of KEGG analysis (Kyoto Encyclopedia of Genes and Genomes) (http://www.kegg.jp), which is a collection of manually curated pathway maps representing hypotheses of the molecular interaction and reaction networks for many biological processes. A subset of differentially expressed genes with a *p*-value < 0.05, FDR < 0.25 and log2 fold change greater or less than +/- 2 was selected and uploaded to STRING and TargetMine for further functional protein association network and gene set enrichment analysis, respectively. Both samples and genes are clustered using Euclidean distances. For genes, an additional elbow function is applied to estimate the number of gene clusters present. Calculated relationships are depicted by dendrograms drawn at the top (samples) and to the left (genes) of the heatmap. The gradation of color is determined by a Z-score that is computed and scaled across rows of genes normalized by TMM.

### Statistical analysis

Each assay was performed with samples of 3 biological replicates (n = 3). All quantitative data were calculated based on results of biological replicates and presented as the mean ± standard deviation. Statistical comparison between groups was analyzed using Student’s t test or one-way ANOVA with a *post hoc* Tukey’s test. A *p*-value of 0.05 was considered statistically significant.

## Acknowledgments

The authors thank Christina Ma, Robert Weishar, To Wang, and Bin Li for assistance with the experimental preparations, Dr. Ellen Leiferman for guiding on animal surgical procedure, and all re-reader including Dr. Ellen Leiferman, En-Lin Hsieh, and Christina Ma.

## Funding

This work was supported by the National Institute of Arthritis and Musculoskeletal and Skin Diseases of the National Institutes of Health under Award Number R01 AR064803. We also thank the University of Wisconsin-Madison Translational Research Initiatives in Pathology Laboratory, supported by the UW Department of Pathology and Laboratory Medicine and UWCCC (P30 CA014520), the Small Animal Imaging & Radiotherapy Facility and Flow Cytometry Laboratory, supported by UWCCC (P30 CA014520) for use of their facilities and services, and the Biotechnology Center’s Bioinformatics Resource Center for assistance with data analysis. The content is solely the responsibility of the authors and does not necessarily represent the official views of the National Institutes of Health.

## Author contributions

M.-S.L. and W.-J.L. designed the experiments; M.-S.L., H.J, H.-C.H., and B.E.W. collected the data; M.-S.L., M.J.S., S.P.P., E.V.S., and W.-J.L. supervised the data collection and analysis; M.-S.L. and W.-J.L. prepared the figures; M.-S.L., H.J, S.P.P., E.V.S., and W.-J.L. interpreted the results; M.-S.L., H.-C.H., and W.-J.L. wrote the article; all authors reviewed the paper.

## Competing interests

The authors declare that they have no competing interests.

## Data and materials availability

All data needed to evaluate the conclusions in the paper are present in the paper and/or the Supplementary Materials. Additional data related to this paper may be requested from the authors.

---

**fig. S1.**
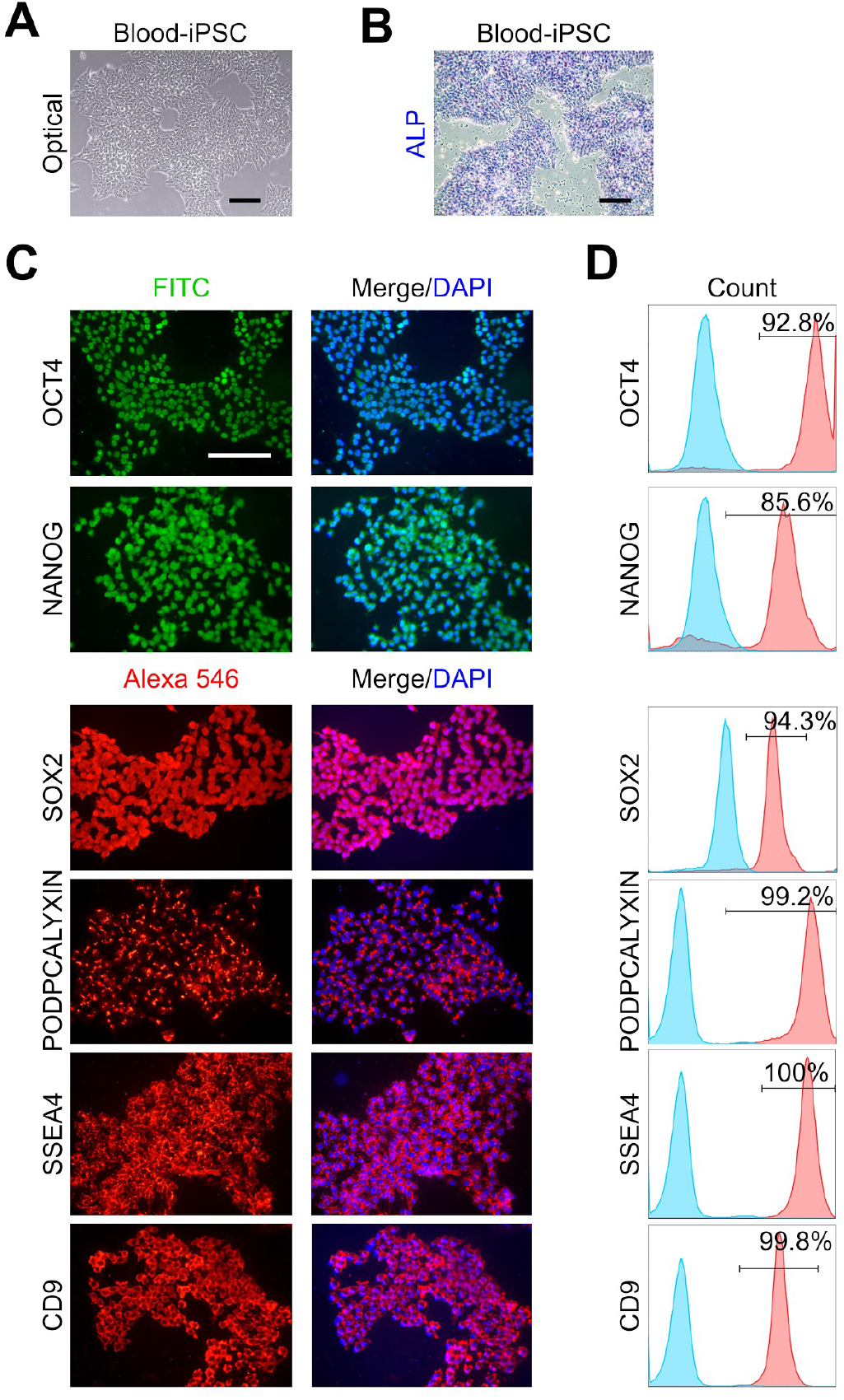
Characterization of blood-derived hiPSCs. **(A)** Morphology of hiPSCs analyzed using bright-field microscopy. **(B)** hiPSCs identified by alkaline phosphatase staining. **(C)** Immunofluorescence staining of hiPSCs for detection of pluripotency markers. Nuclear DNA was labeled with DAPI. **(D)** Percentages of hiPSCs stained positive for corresponding pluripotency-related markers determined by flow cytometry. *n* = 3. Scale bar, 200 μm.

**fig. S2.**
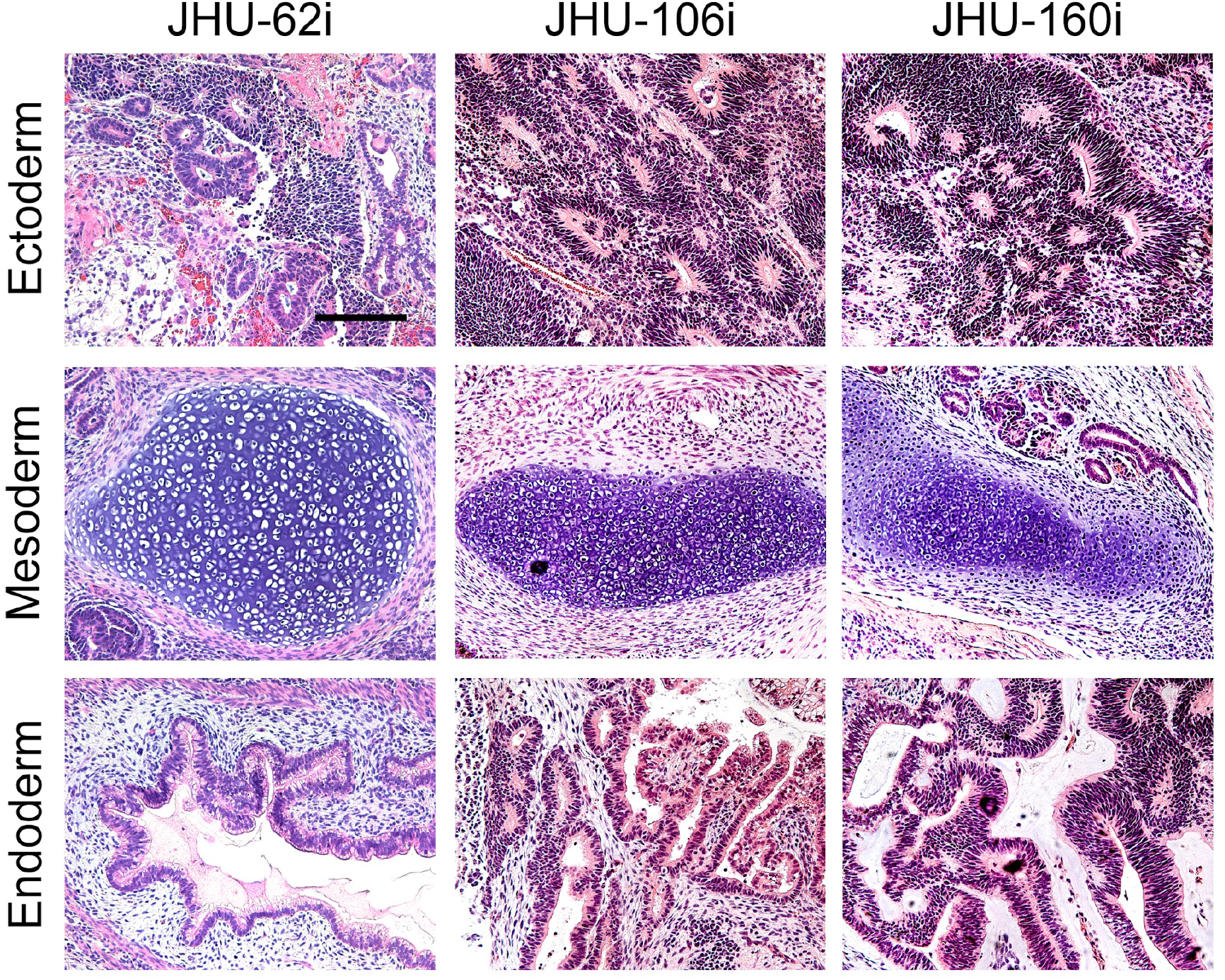
Teratoma assay for evaluation of hiPSC pluripotency. Teratoma formation in mice receiving three independent blood-derived hiPSC lines analyzed by H&E staining. Ectoderm is represented by neural epithelium, mesoderm by cartilage, and endoderm by gut. *n* = 3. Scale bar, 200 μm.

**fig. S3.**
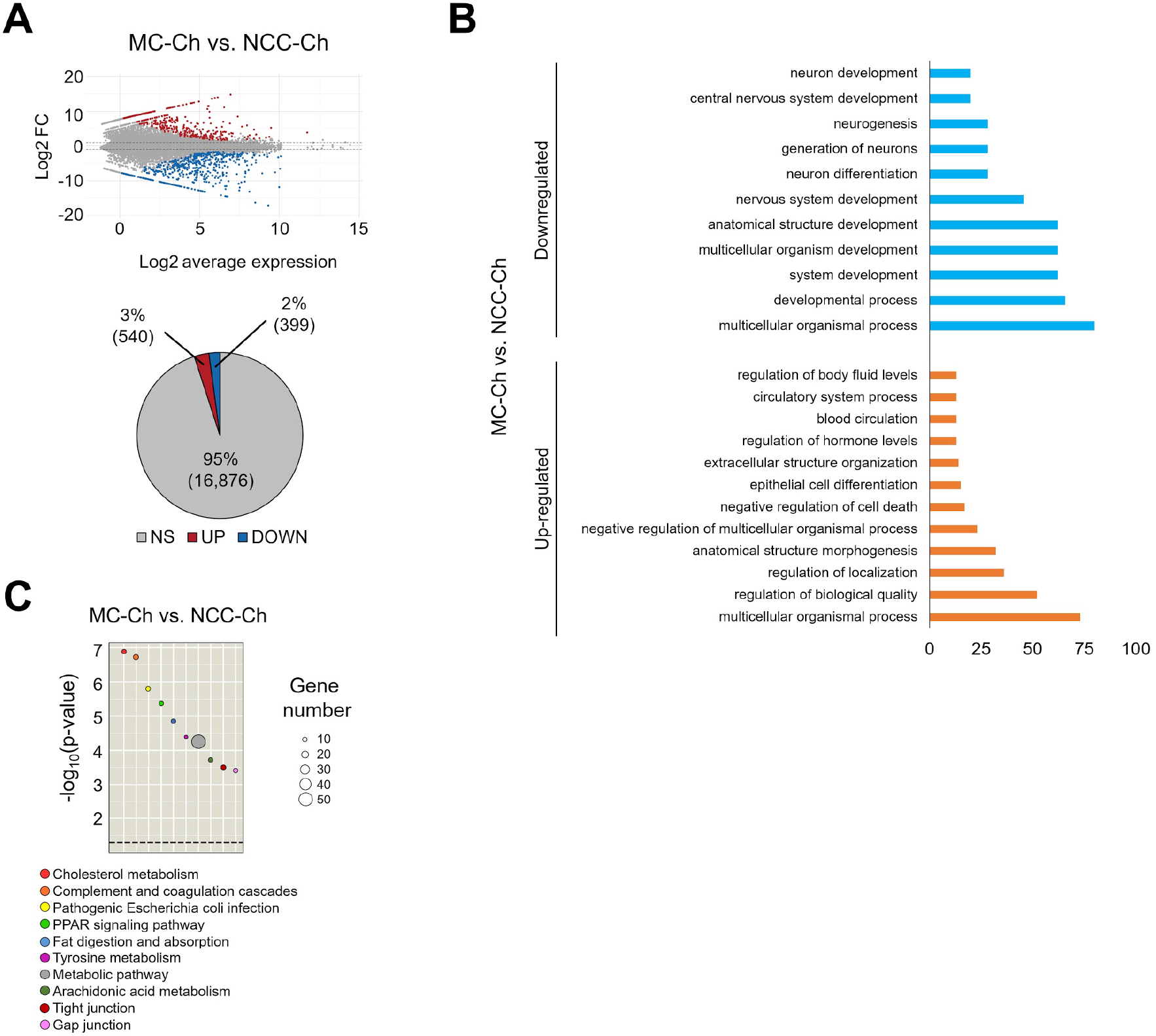
Comparison of global transcriptome profiles between MC-Chs and NCC-Chs. **(A)** Smearplot presenting differences in global transcript expression of NCC-Chs compared to MC-Chs. Red and blue dots denote individual up- and down-regulated genes, respectively, at the significance threshold of adjusted *p*-value of 0.05. Gray dots reflect those genes with no statistically significant differential expression. Percentages of the up-regulated (Up), down-regulated (Down), and non-significant (NS) genes are shown in corresponding pie charts. **(B)** Gene ontology enrichment analysis of MC-Chs compared to NCC-Chs. Categories of the most differentially expressed gene sets with a *p*-value less than 0.05 are shown. The number of genes in each enrichment category is represented by the length of an individual bar. **(C)** Analysis of KEGG pathway enrichment identifying abundance of differentially expressed genes in predominant biological categories of MC-Chs compared to NCC-Chs. Significantly enriched KEGG pathways are ordered from most to least significant. The number of genes in a specified KEGG pathway is denoted by the size of a circle. The color of each circle corresponds to that of the specific KEGG pathway. The dashed line represents significance at *p*-value = 0.05. *n* = 3.

**fig. S4.**
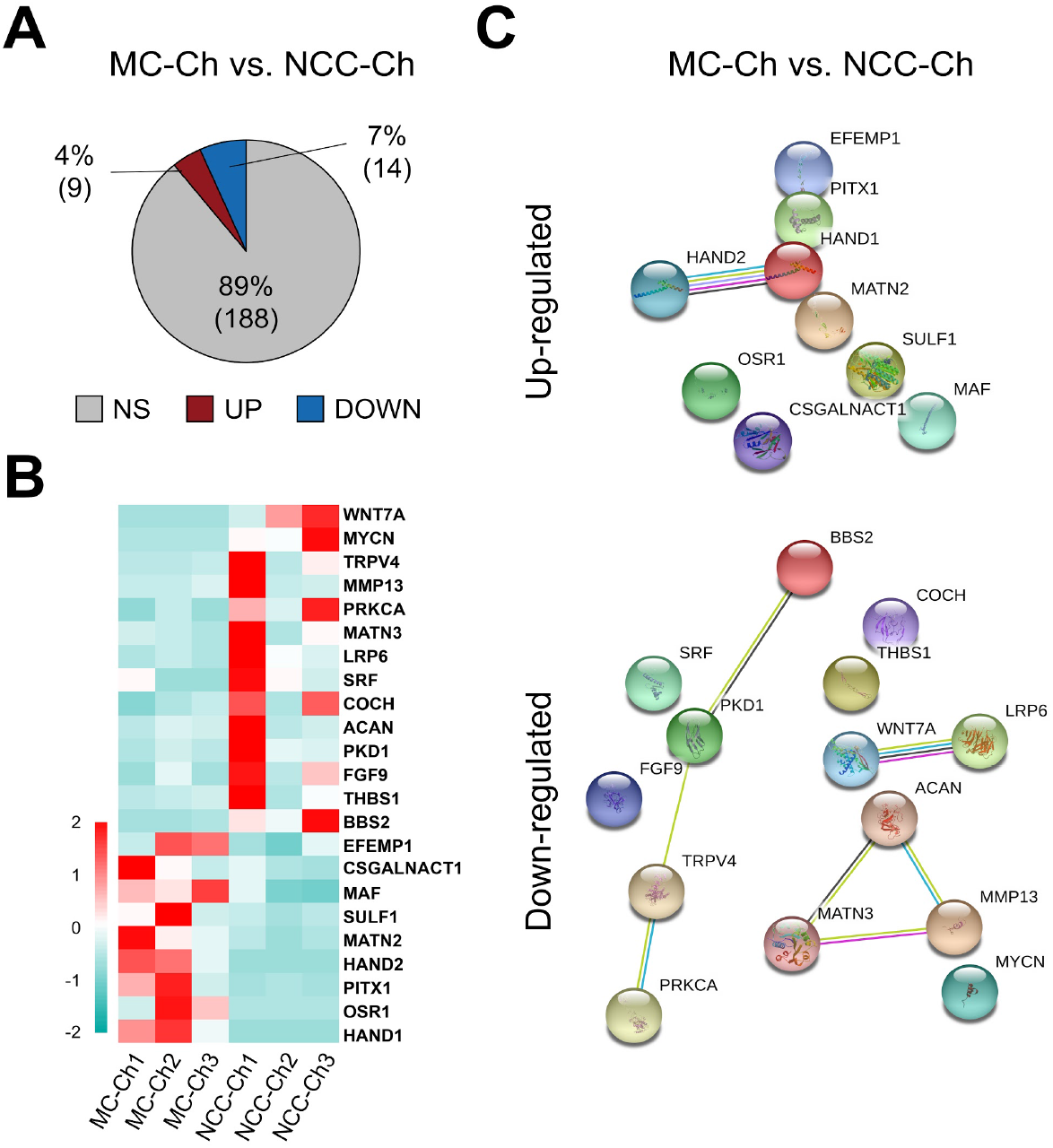
Analysis of differentially expressed transcripts associated with cartilage development between MC-Chs and NCC-Chs. **(A)** Pie chart depicting percentages of the up-regulated (Up), down-regulated (Down), and non-significant (NS) transcripts in MC-Chs compared to NCC-Chs. **(B)** Heatmaps of differentially expressed cartilage development-associated transcripts between MC-Chs and NCC-Chs. The color intensity of each grid denotes the extent of change in transcript expression, where up-regulated transcripts are shown in red and down-regulated ones in turquoise. **(C)** Functional protein association networks established using the STRING database showed known and predicted interactions of cartilage development-associated molecules differentially expressed between MC-Chs and NCC-Chs. *n* = 3.

**fig. S5.**
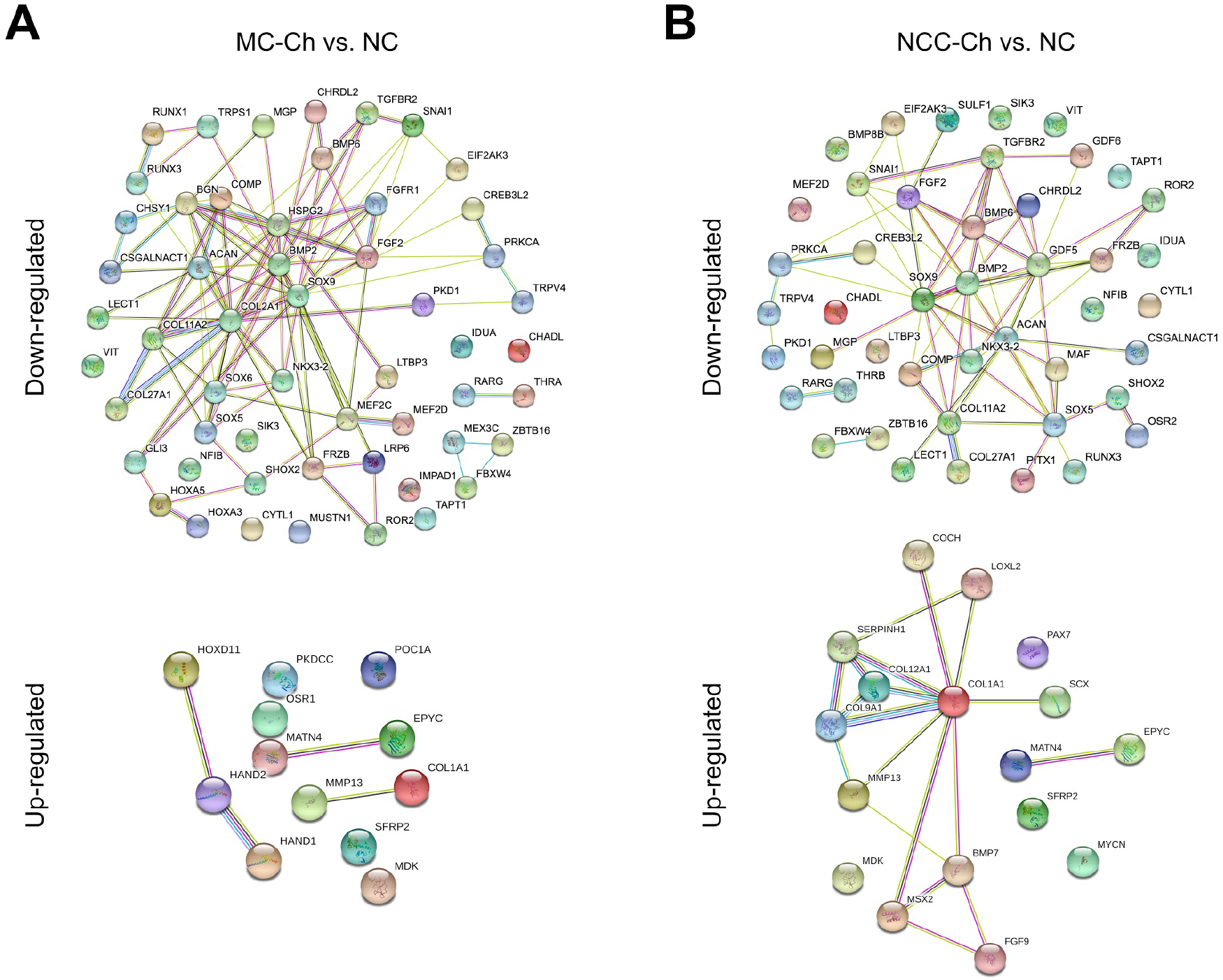
Interactions of differentially expressed cartilage development-associated molecules in hiPSC-derived and native chondrocytes. Functional protein association networks established using the STRING database showed known and predicted interactions of cartilage development-associated molecules differentially expressed between MC-Chs **(A)** or NCC-Chs **(B)** and NCs. *n* = 3.

## Notes

### Competing Interest Statement

The authors have declared no competing interest.

